# Spatial predictive context speeds up visual search by biasing local attentional competition

**DOI:** 10.1101/2023.07.14.548976

**Authors:** Floortje G. Bouwkamp, Floris P. de Lange, Eelke Spaak

## Abstract

The human visual system is equipped to rapidly and implicitly learn and exploit the statistical regularities in our environment. Within visual search, contextual cueing demonstrates how implicit knowledge of scenes can improve search performance. This is commonly interpreted as spatial context in the scenes becoming predictive of the target location, which leads to a more efficient guidance of attention during search. However, what drives this enhanced guidance is unknown. First, it is under debate whether the entire scene (global context) or more local context drives this phenomenon. Second, it is unclear how exactly improved attentional guidance is enabled by target enhancement and distractor suppression. In the present MEG experiment, we leveraged Rapid Invisible Frequency Tagging (RIFT) to answer these two outstanding questions. We found that the improved performance when searching implicitly familiar scenes was accompanied by a stronger neural representation of the target stimulus, at the cost specifically of those distractors directly surrounding the target. Crucially, this biasing of local attentional competition was behaviorally relevant when searching familiar scenes. Taken together, we conclude that implicitly learned spatial predictive context improves how we search our environment by sharpening the attentional field.

## Introduction

Locating a relevant item in a crowded visual field, also known as *visual search*, is a critical skill guiding our behavior in daily life. Prior experience yields a vast amount of knowledge on how items tend to co-occur in scenes. Knowledge of these relations can narrow down the search space by generating expectations about *what* to expect, and *where* to expect it (Peelen et al., 2023; Võ et al., 2019). The implicit and automatic learning of regularities is called *statistical learning.* This is a very useful tool in investigating how we learn from our environment, and how we exploit that knowledge in our interaction with the world.

Within the domain of visual search, a classic example of statistical learning is *contextual cueing*. Chun & Jiang (1998) discovered that participants are markedly faster in finding a target in search scenes that are repeated, compared to search scenes that are novel. After (implicit) learning, the predictive relationship between spatial context and target location is exploited, leading to more efficient guidance of spatial attention when searching these scenes (Goujon et al., 2015; Y. V. Jiang et al., 2019; Sisk et al., 2019). This improved attentional guidance can be interpreted as the result of an optimized attentional priority map (Sisk et al., 2019; Wolfe, 2021), but how exactly this learned spatial context alters what we prioritize, is an open question.

Firstly, there is debate on the extent of context involved in contextual cueing. Some have argued that only *local* context directly surrounding the target is learned, since repeating only such local context produced a similar effect to classical contextual cueing (Brady & Chun, 2007). However, this does not hold when presentation time is limited (Xie et al., 2020), and there has been evidence that distractor context *unrelated* to the target is also learned (Beesley et al., 2015a). The question thus remains whether local and/or global context is learned in contextual cueing (Goujon et al., 2015).

This question resides in the larger debate of if and how chunks are learned from patterns that have statistical co-occurrences. Similarly to the notion of chunk extraction, (Conway, 2020; Perruchet, 2019), a configuration is learned rather than element wise associations. It is, however, unknown whether all predictive context is automatically used in this configural learning, or, alternatively, if the interaction between attention and statistical learning (Conway, 2020) leads to a focus on local context only.

Secondly, more efficient guidance of selective attention can be achieved by either enhancing the target, or suppressing distractors, and there is behavioral evidence implicating both (Makovski & Jiang, 2010; Ogawa et al., 2007). Additionally it has been convincingly shown that not only distractor-target relationships are learned in contextual cueing. Instead, only repeating the distractors also leads to contextual facilitation, albeit a smaller effect (Beesley et al., 2015b; Vadillo et al., 2021). This implies that context learning can lead to distractor suppression, independent of the target. However, when the context *is* predictive of the target, both distractor suppression and target enhancement are potential mechanisms responsible for improved attentional guidance. It remains unclear, what the relative contributions and the neural consequences are of these two mechanisms. This is a relevant question, as target enhancement and distractor suppression are thought to rely on distinct (neural) mechanisms (Noonan et al., 2016; Slagter et al., 2016; Wöstmann et al., 2022).

We set out to investigate how spatial predictive context modulates attentional guidance during visual search at the stimulus level: does it involve mainly local or global context? And what are the neural consequences of spatial predictive context learning? More specifically, what is the relative contribution of target enhancement and distractor suppression to activity in early visual regions? For this, we leveraged Rapid Invisible Frequency Tagging (RIFT) (Drijvers et al., 2021; Minarik et al., 2022; Seijdel et al., 2023; Zhigalov et al., 2019). Visual stimulation by manipulating the luminance of a stimulus periodically will elicit an electrophysiological signal that has the same frequency as the stimulation. This steady-state visual evoked potential (SSVEP) can be detected with non-invasive electrophysiological methods, such as electroencephalography (EEG) and magnetoencephalography (MEG). The distinct advantage of *rapid* tagging is that it is invisible to the human eye, and therefore does not interfere with perceptual processes, while signal-to-noise ratio in neural (EEG/MEG) recordings is still very high (Minarik et al., 2022). Importantly, the magnitude of the RIFT signal is modulated by selective attention (Zhigalov et al., 2019). This enables us to measure both stimulus processing, and the attentional modulation thereof, with high temporal and stimulus precision.

By tagging three stimulus types with unique frequencies we were able to track the attentional processing of the target, a distractor in the local context of the target and a distractor further from the target, i.e. the global context. We hypothesized that the advantage after contextual learning will be accompanied by target enhancement and distractor suppression at the level of early visual cortex, to which we should be sensitive using RIFT. If only local context is involved, we expected to find neural modulation of only the distractor near the target. If, however, global context is encoded, we expected neural modulation of both the near and far standing distractor. Additionally, if target enhancement and distractor suppression operate independently, we anticipated a distinct temporal profile of distractor suppression preceding target enhancement.

In brief, we successfully tagged multiple stimuli and were able to track online attentional processing of both target and distractors, demonstrating the strength of RIFT even in visual competitive settings such as visual search. Behaviorally, we found improved visual search of repeated compared to novel scenes. Neurally, we found simultaneous target enhancement and suppression of distractors near the target, effectively biasing local attentional competition. Specifically this biasing of local attentional competition improved behavioral performance when searching repeated scenes. We conclude that spatial predictive context enhances attentional guidance by sharpening the attentional field.

## Methods and materials

### Data and code availability

All data and code used for stimulus presentation and analysis are freely available on the Donders Repository. The link to the repository will be accessible to reviewers and be publicly available upon publication at https://doi.org/10.34973/xxnc-4x21.

### Participants

Thirty-six adults (23 identifying as female and 13 identifying as male, age 18-49 years, M = 28.95, SD = 8.72) participated in the experiment. All were recruited via the Radboud University participant database and informed consent was given before the start of the experiment. The study was approved by the local ethics committee (CMO 2014/288; CMO Arnhem-Nijmegen, the Netherlands) and was conducted in compliance with these guidelines. None reported a history of neurophysiological disorders and all had normal or corrected-to-normal vision. We aimed for a sample size of 34 participants, based on *a priori* power analysis (G*Power 3.1)(Faul et al., 2009) computing the required sample size to achieve a power of 0.8 to detect a medium effect size of Cohen’s *d* = 0.5, at α = 0.05, for a two-tailed paired *t* test. Four participants were excluded due to insufficient data quality (see MEG analyses). The final sample therefore consisted of 32 participants.

### Stimulus material

Search scenes consisted of scenes with 8 stimuli: one target letter T and seven distractor L shapes, with a small (10%) offset in the line junction to increase search difficulty (Y. Jiang & Chun, 2001). Stimuli measured 2.5° x 2.5° and were displayed as mid grey on a dark grey background. Distractor shapes were rotated randomly with a multiple of 90°. The target was tilted either to the left or to the right. Each scene had a fixation dot at the center (outer white diameter 8 pixels, inner white diameter 4 pixels). Stimuli were placed on a 7x5 grid spanning the screen from - 11° to 11° horizontally and -7.5° to 7.5° vertically, excluding the center position. To prevent collinearity the stimuli were jittered with ± 0.5°(Chun & Jiang, 1998b). To ensure search scenes were approximately equal in difficulty, we generated search scenes (both Old and New) abiding multiple restrictions. The target was always placed between 6° and 10° of eccentricity, the mean distance between target and distractors was kept between 9° and 11°, no more than one quadrant was empty, and in quadrants with >2 stimuli, local crowding was prevented. Additionally, to prevent location probability learning, target locations across the repeated, Old, trials were evenly distributed over the four quadrants, meaning there were no target-rich versus target-sparse quadrants across search scenes (Y. V. Jiang et al., 2013). Three types of stimuli within these scenes were tagged with unique frequencies (see Frequency tagging): the target, a distractor near (D*near*) the target and a distractor further (D*far*) from the target. The far standing distractor was always placed at the opposite position of the target relative to the center of the screen. The distractor closest to the target was selected as Dnear, with a minimum distance of *less* than 4° from the target, while being minimally 4° from the center of the screen.

### Experimental conditions

Each block consisted of 32 trials and half of the trials each block were newly created (*New)*, while the other half of trials were always the same 16 search scenes, rendering them *Old* from block 2 onwards (figure 1B). In these Old displays, both (jittered) location and orientation of distractors were repeated, as well as target location. Target orientation was always randomized (tilted left or right) to prevent any learning of the correct response in Old trials. Old and New trials were presented in pseudo-randomized order. Practice trials consisted of a separate set of scenes without any repetitions.

**Figure 1.**
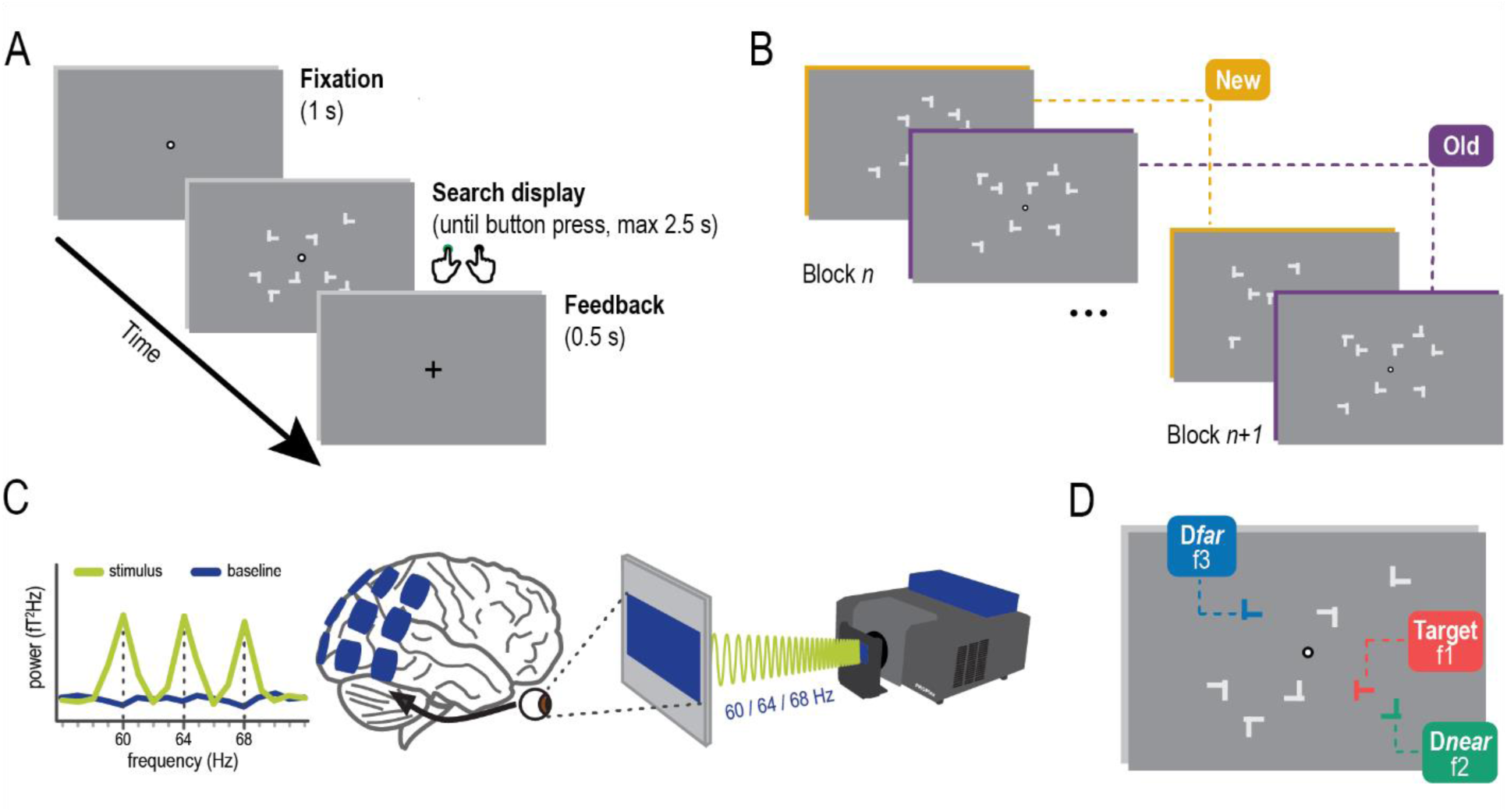
Paradigm. A) Task. Participants engaged in a visual search task where they had to locate a target letter T embedded amongst distractor L-shapes and report its orientation with a left or right button press. B) Manipulation. Half of all trials consisted of search scenes that were repeated every block, rendering them ‘Old’. The other half of the trials consisted of unique and newly created search scenes. C) RIFT. The projector allows rapid tagging of stimuli at our frequencies of interest. This tagging signal can be identified in the power spectrum as sharp peaks at the frequencies used (image adapted from Seijdel et al 2023) D) Tagging search scenes. Within each search scene three types of stimuli were tagged at a unique frequency: the target, a near-standing distractor and a far-standing distractor. Assignment of frequencies to stimuli was counterbalanced across participants. Colors of the different stimulus types are for illustration purposes only.

### Frequency tagging

To achieve Rapid Invisible Frequency Tagging, we used a PROPixx DLP LED projector (VPixx Technologies Inc., Daint-Bruno-de-Montarville, Canada). This monitor can interpret four quadrants of the screen in three color channels of the GPU screen buffer, as twelve separate smaller grayscale frames, which it then projects in rapid succession. This leads to a twelvefold increase of the 120 Hz presentation rate, yielding 1440 Hz. Three stimuli (figure 1C) were tagged with either 60, 64 or 68 Hz, and stimulus-frequency mapping was counterbalanced across subjects. Color values of the stimuli oscillated between 0 (black) and 255 (white), at the assigned frequency by multiplying the max luminance value with a sinusoid of the relevant frequency. Other stimuli were kept constant at the mean value of 127. All stimuli therefore appeared as mid grey on the screen. To prevent straining the eyes, the background was kept at dark grey (value = 43). No frequency tagging was applied in the final two blocks, and all stimuli were kept constant at the mean value of 127. These trials served as a baseline to ensure that power differences were due to the frequency tagging.

### Experimental procedure

After verbal instructions and signing informed consent, participants were seated in the dimly-lit magnetically shielded MEG room wearing a neck brace for head stabilization. Stimuli were presented on a screen at approximately 85 cm distance, using Matlab 2018b (Mathworks Inc, Natrick, USA) and the Psychophysics Toolbox, (Brainard, 1997; Pelli, 1997). The experiment started with a 9 point calibration session for the eye tracker, which was used to monitor fixation of the participant throughout the experiment. Participants were introduced to the task and instructed to report the orientation of the Target (either tilted left or right) with a left or right button press.

Furthermore, it was stressed that participants had to remain their fixation at the center of the screen. Each trial (figure 1A) started with a fixation period of one second, followed by the search scene until the subject pressed a button to indicate their response. Maximum response time was 2.5 seconds. At the end of each trial they received feedback (0.5 seconds) on performance with a symbol: V for correct, X for incorrect and O for being too late. Participants practiced this task for 64 trials. If either performance or fixation was insufficient, they practiced for another 64 trials. Then, the main experiment started, consisting of 26 blocks offered in sets of two blocks of 32 trials each. In between these ‘subject blocks’ of 64 trials, participants received feedback on performance (% correct and average response time in seconds) and were offered a short break. When ready for the next block, their head position was realigned if necessary before starting. At the end of the main experiment, participants performed a short recognition task where they had to judge for 32 search scenes (the 16 Old scenes and 16 Newly generated scenes) whether this scene was familiar or not. Responses in the recognition block were self-paced (no timeouts). The entire experiment lasted approximately 1.5 hours.

### Behavioral analyses

Analyses and visualization of behavioral data was done using R (R Core Team, n.d.) and packages ggplot2 (Wickham, 2016), raincloud plots (Allen et al., 2021), Effsize (Torchiano, Marco, 2016) and BayesFactor (Morey & Rouder, 2022) for R. Reaction time was our primary, and accuracy our secondary variable of interest. Only trials were a response was given in time were included in the analyses (only 1 subject was too late on 2 trials), trials with an response given before 100 ms were regarded as accidental presses and excluded (3.85% of trials). Reaction time analyses were performed on correct trials only. We improved normality of the reaction times by log10-transforming the raw values. Statistical tests are based on these log-transformed reaction times; however, we report and plot raw reaction times in the Results section for interpretability. Plots over experiment time of both reaction times and accuracy are smoothed across neighboring blocks. Statistical assessment focused on the latter part of the experiment (blocks 9-26, unsmoothed, selected a priori) when learning is assumed to have taken place, which was assessed with an analysis of variance. We then directly contrasted conditions by means of a paired samples t-test. Accuracy on the recognition task at the end of the experiment was pitted against the null hypothesis of random guessing (50% accuracy) with a one sample t-test. To assess the size of any effects we report general eta-squared for F-tests and Cohen’s d for T-tests. In addition, we quantified the relative evidence for the alternative hypothesis against the null hypothesis using Bayes factors.

### MEG acquisition

During the main experiment, MEG activity was recorded using a 275-channel axial gradiometer CTF MEG system (CTF MEG systems, Coquitlam, Canada) situated in a magnetically shielded room (MSR). An online 300hz low pass filter was used and data was digitized with a sampling rate of 1200 Hz. The head position of participants was monitored in real time using markers on the nasion, and left and right periauricular points (Stolk et al., 2013). Participants were asked to readjust their head position when the deviation from the original starting point was larger than 5 mm. Additionally, participants’ gaze was monitored using an SR Research Eyelink 1000 eye tracker and responses were collected using an MEG-compatible button box.

### MEG preprocessing

MEG data was preprocessed and analyzed using the Fieldtrip toolbox (Oostenveld et al., 2011) in a Matlab environment (Mathworks, version 2018b). The data was segmented into trials and preprocessed by first applying third order gradient denoising and general demeaning of the data. Subsequently, trials affected by high noise or muscle movement were removed using a semi-automated method by visually identifying unique high variance trials. Then, the data was down-sampled to 400 Hz and cleaned of any residual artifacts due to eye movements, heartbeat or other sources of noise using Independent Component Analysis (Bell & Sejnowski, 1995; Jung et al., 2000). During recording, the MEG system was affected by an occasional drifting high-frequency artifact caused by external RF-noise entering the MSR. This artifact was detected in six of the 36 acquired datasets, and in three of these it drifted through our frequency range of interest during a subset of trials. All six datasets were successfully cleaned in the preprocessing stage with an in-house procedure (Principal Component Analysis applied with the artifact as a reference signal). Together, these preprocessing procedures led to the removal of on average 6.48% of trials.

### MEG analyses

For the assessment of general tagging we used all tagged trials, independent of condition. We discarded trials that were shorter than 800 ms. This choice was made after processing, but before the analyses. We needed to consider the trade-off between sufficient length to analyze a meaningful time window reflecting the search process, and not cutting too many trials (800 ms: 15.5 % of trials on average, 49.94% of discarded trials were Old). Data was then cut to an a priori defined time window of interest (0.2 – 0.8 s post stimulus onset) and a baseline window (- 0.5 – 0 s pre stimulus onset). As tagged stimuli varied in location per search scene, we applied trial-based spectral analyses. Tagging power (i.e., the magnitude of the neural signal corresponding to the stimulus-specific tagging frequency), was estimated using a boxcar window focused on the narrow band responses at our three tagging frequencies of interest. Data was zero-padded to 1 second in order to obtain a frequency spacing of 1 Hz. We approximated planar gradiometer data by converting our axial gradiometer data to orthogonal planar gradiometer pairs, and computing power separately before combining them by averaging. This enables a more straightforward interpretation of the MEG data, as planar gradient maxima are thought to be centrally located above their neuronal source (Bastiaansen & Knösche, 2000). Power during the window of interest was expressed as decibels relative to the baseline period. Based on this data, we selected the twenty sensors within occipital-parietal region that were most responsive to average power at the tagged frequencies per individual subject (results section, figure 3B). We then inspected the condition-averaged power spectrum (frequency-resolved from 52 to 76 Hz) at the selected sensors per subject. This led to the exclusion of 4 subjects that had very noisy power spectra, with no clear peaks at the frequencies of interest and/or additional peaks at neighboring frequencies. This yielded our final sample size of 32 subjects. For our main analyses we subsequently selected only correct (target orientation task) trials from the latter part of the experiment (a priori decision, latter 2/3 of all tagged blocks), and considered the initial 8 blocks learning blocks (Chun & Jiang, 1998b). We separated the trials into our Old and New condition.

Because trials varied in length we analyzed the data both aligned to stimulus onset (stimulus-locked) and aligned to the response (response-locked). We subjected the data to spectral analyses, both averaged over a priori defined time windows of interest (0.2 - 0.8 s post-stimulus or –0.6 - 0 pre-response) following the procedure described above, and in a time-resolved manner (–0.5 - 0.8 s stimulus-locked or –0.8 – 0.5 response-locked). Time-resolved power was estimated using a 250 ms sliding window (min. 15 cycles) in steps of 50 ms, after zero-padding the data to 2 seconds. Baseline normalization was done within condition. Statistical analyses of the power values per stimulus type were done by contrasting the target to both distractor types, with a one-tailed paired samples t-test, testing for the increase in power of target over distractors. Additionally we tested the difference between the two distractor types with a two-tailed paired samples t-test. Statistical analyses on the time-resolved power were done with non-parametric cluster based permutation tests (Maris & Oostenveld, 2007). For this, only the time windows post-stimulus and pre-response, taking temporal leakage into account, were considered (0.15 - 0.6 s stimulus-locked or –0.6 – 0.15 response-locked). Clustering threshold was set to alpha = 0.1, the number of randomizations was set to 10,000, and the test statistic was based on a paired samples t-test. Again we contrasted target to both distractors using a one-tailed test, while contrasting the distractors using a two-tailed test.

To assess behavioral relevance, we tested whether a larger difference in power between stimuli was related to faster reaction times. For this, we used the trial-based stimulus-locked power of Old and New trials, since only stimulus-locked power will yield trial-by-trial variance that can be related to variance in reaction time. We correlated the difference between target and each distractor to the behavior (log-transformed reaction times) on these trials within subject. To specifically isolate effects related to the learning of Old scenes (rather than effects that may be general to any visual search process), behavioral relevance was assessed not only across all trials, but also specifically across the 16 displays within the Old condition. For these analyses, both power and reaction times on Old trials were averaged across trials per display (16 in total). We then, for both analyses, computed the Pearson correlation values across displays, per subject, and tested the distribution of these correlations over subjects against null with a one-sided one sample t-test. In addition to Cohen’s d to assess effect size, we quantified the weight of evidence by means of Bayes Factors. Bayes Factors were either one sided or two sided, matching the frequentistic statistics (Keysers et al., 2020).

## Results

Participants were seated in the MEG while engaged in a visual search task. As typical for contextual cueing, half of all trials were search scenes randomly generated each block, labeled as ‘New’. The other half were search scenes that were repeated every block, which we labeled as ‘Old’. To assess the mechanisms underlying enhanced attentional guidance by spatial predictive context in Old trials compared to New trials, we tagged the Target and two distractors in each search display using Rapid Invisible Frequency Tagging (RIFT). By tagging both a distractor *near* the target (D*near*) and a distractor *far* from the Target (D*far*), we were able to investigate the roles of local and global context when spatial context becomes predictive of target location.

### Improved search performance on Old trials compared to New trials

For all analyses, both behavioral and MEG, we focused on the latter two thirds of the experiment (blocks 9 and onward, selected a priori), after participants putatively learned the predictive spatial context within Old trials.

#### Accuracy

Overall accuracy on the task was high (Mean 85.90% ± 6.08% SD; range: 68.78% - 95.87%) and it improved over time (first/latter part: F_1,31_ = 27.27, p < .001, η^2^_G_ = .10; Figure 2A). There was, however, no difference between Old and New scenes in accuracy (condition: F_1,31_ = .31, p = .31, η^2^_G_ = .002, condition x time: F_1,31_ = 2.70, p = 0.11, η^2^_G_ = .004; Figure 2B), allowing us to focus solely on reaction times.

**Figure 2.**
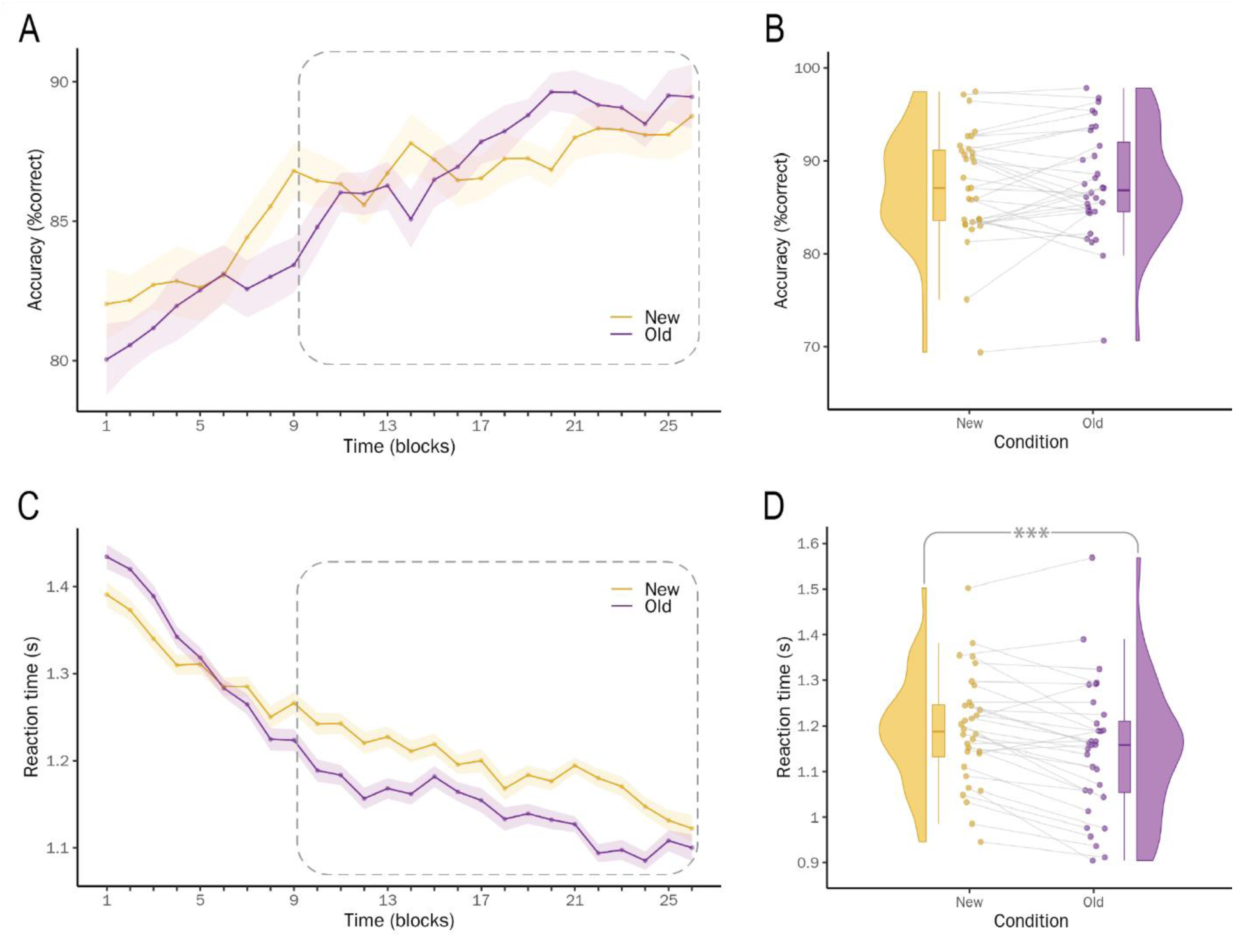
Behavioral results. Accuracy. (A) Percentage correct (smoothed across neighboring blocks, by taking the mean of block N, N − 1 and N + 1) plotted over the time course of the experiment (shading indicates within-participant corrected standard error of the mean). Dotted box indicates the trials used contrasting Old to New scenes, when contextual learning is assumed to have taken place. (B) Accuracy on New versus Old trials on learned trials. Dots are individual participants. The boxplot indicates the interquartile range (IQR, the box), the median (bar on box) and the minimum and maximum value within ±1.5 x IQR (whiskers). (C) Reaction time plotted over the time course of the experiment (D) Reaction times on New versus Old trials on learned trials.

#### Reaction times

As the distribution of reaction times is typically non-normal, all analyses are based on log-transformed data. For interpretability, we will report raw RT values in the text. Overall search speed was well within the time limit of 2.5 seconds (1.21 s ± 0.43 s). Participants became faster in in general (time: F_1,31_ = 5.31, p < .001, η^2^_G_ = .23). Additionally, there was a difference between Old and New scenes (condition: F_1,31_ = 56.32, p = .028, η^2^_G_ = .004) that interacted with time, revealing learning (time x condition: F_1,31_ = 27.69, p < .001, η^2^_G_ = .014). Focusing on the latter part of the experiment, participants were faster in finding the target on Old trials compared to New scenes (New: 1.19 s ± 0.41 s and Old: 1.14 s ± 0.41 s, difference: 50 ms ± 60 ms, t(31) = 4.87, p < .001, *d* = .33, BF_10_ = 733) And these results also hold when not normalizing the reaction times (t(31) = 4.49, p < .001, *d* = .32, BF_10_ = 280). We thereby replicated the classic contextual cueing effect (Figure 2C and D). Even though targets were evenly distributed across quadrants, one may wonder whether participants might have learned the marginal distribution of target locations in Old scenes, rather than the relationship between context and target location. To examine this possibility, we tested whether New trials with targets at exactly the same locations as Old trials (but no spatial predictive context) resulted in faster reaction times compared to New trials where the target was elsewhere. The data compellingly suggest that this was not the case (BF10 = 0.02). The reaction time advantage for Old trials is thus due specifically to the presence of spatial predictive context.

### Identifiable power peaks at all tagged frequencies

As a first step in analyzing the MEG data, we calculated power at our frequencies of interest pooled over conditions and participants. When normalizing power with a baseline period and plotting the spectra (Figure 3A), we observed clear peaks at the tagged frequencies of interest, with no apparent differences in power between the three frequencies. Additionally, we observed no such peaks for trials where the search scenes were not tagged (Fig 3A, blue line). This steady state visual evoked response was, as expected, localized at posterior sensors, covering primary visual cortex (Figure 3B). We can therefore conclude that RIFT can be used to reliably tag several relatively small (∼2.5 degrees) non-foveated stimuli simultaneously on the screen, at variable locations in retinotopic visual space.

**Figure 3.**
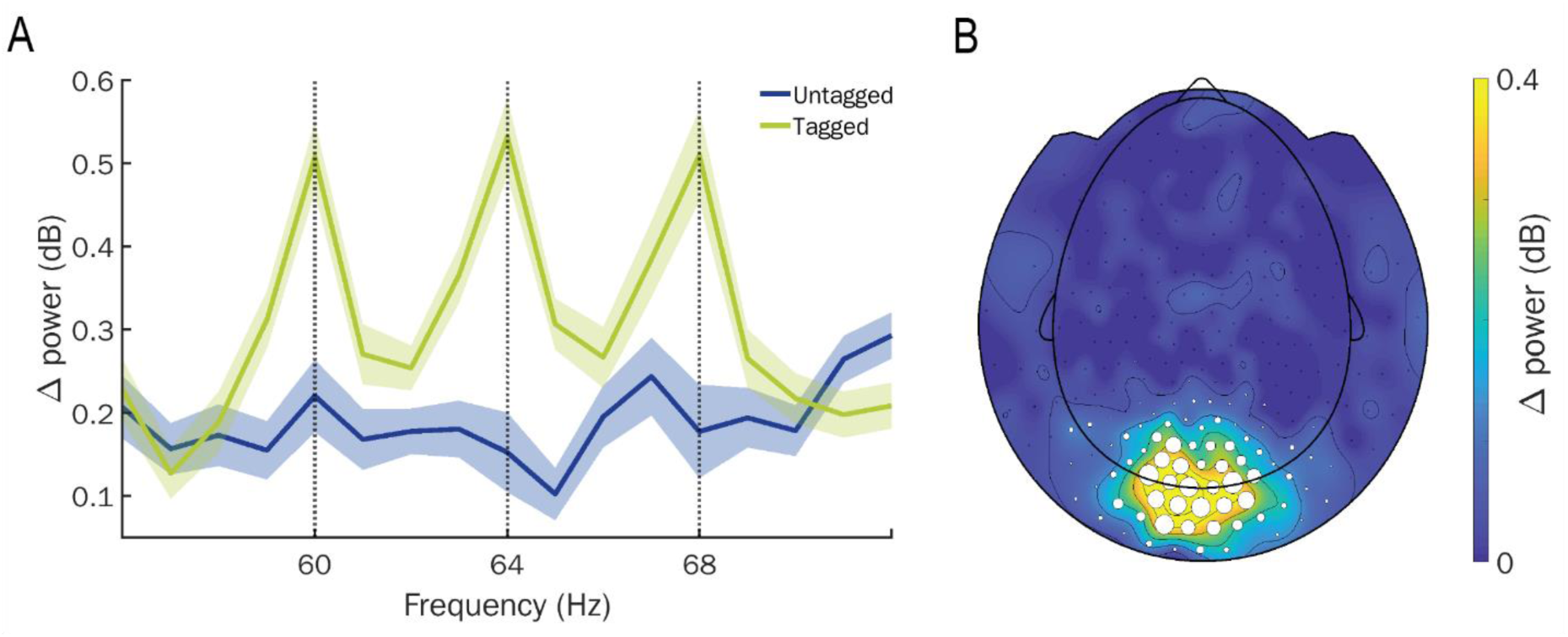
Power spectrum and topography of RIFT. A) The power spectrum shows clear peaks at the frequencies of interest, which are absent on trials of the last two blocks when the search scenes were not tagged. The topography of (planar) power at the frequencies of interest indicates that tagging power is mostly seen in the posterior occipital region. White circles indicate sensors that were selected for the analyses, with the size indicating for how many participants a given sensor was used.

### Stronger power for target compared to distractors when searching Old scenes

To ensure that quality of fixation is not weighing in on our RIFT results, we checked whether variance in eye position differed between conditions. An unsigned t-test comparing the mean variance in x and y direction across the trials we used for data analyses revealed no difference between Old and New : t(31) = .89, *p* = .38, *d* = .005, BF_10_ = .27.

For our main analyses, we calculated power at the tagged frequencies separately for trials where participants were searching Old and New scenes. Since visual search trials vary in length, we analyzed these trials both locked to stimulus onset and locked to the response. We analyzed tagging power values both averaged over a priori defined time windows of interest (0.2 to 0.8 s post-stimulus, and −0.8 to −0.2 s pre-response), as well as in a time-resolved manner using cluster-based permutation tests. As we are specifically interested in attentional competition between the tagged stimuli during visual search, we contrasted target power with power for both types of distractors (D*far* and D*nea*r) and additionally tested the difference between local and global context (D*nea*r versus D*far*).

First, we looked at regular visual search trials without spatial predictive context (New trials). We found no statistically significant differences for any of our comparisons within the a priori time window analysis (Figure 4A): neither between Target and distractors (Target > D*near* stimulus-locked t(31) = 1.12, *p* = .14, *d* = .12, BF_10_ = .57 and response-locked t(31) = .82, *p* =.21, *d* = .08, BF_10_ = .40; Target > D*far* stimulus-locked t(31) = .14, *p* = .44, *d* = .01, BF_10_ = .21 and response-locked t(31) = .09, *p* = .46, *d* = .01, BF_10_ = .20) nor between distractors (D*near* <> D*far* stimulus-locked t(31) = 1.15, *p* = .26, *d* = .11, BF_10_ = .34 and response-locked t(31) = .84, *p* = .41, *d* = .08, BF_10_ = .26). Time-resolved results also did not reveal any significant cluster-based differences between the stimulus types during New trials (Figure 4B), neither stimulus-locked nor response-locked.

**Figure 4.**
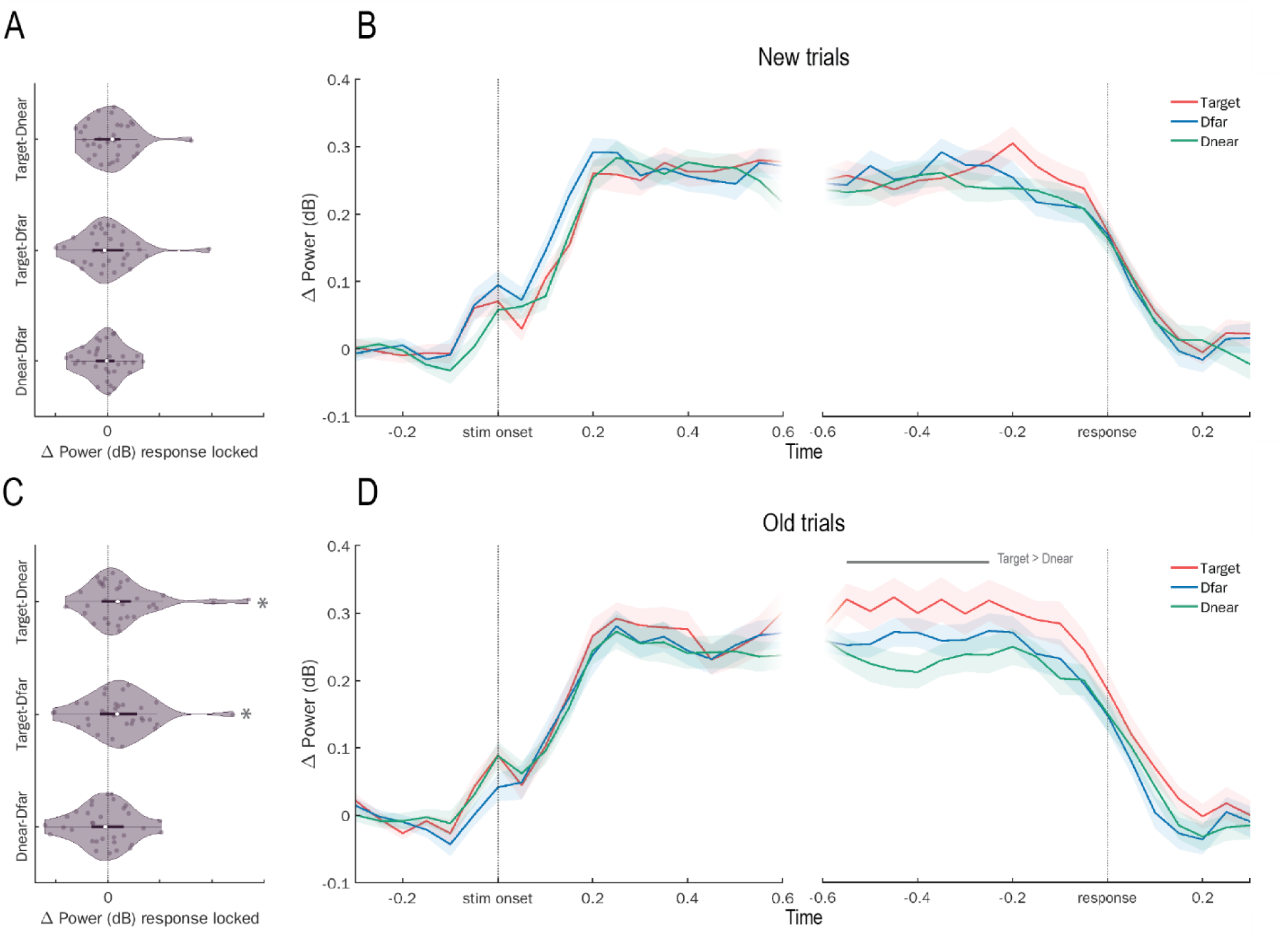
Power Old and New trials. (A) Response-locked power New trials averaged over a priori time window (−0.6 to −0.2 s before response), contrasting the stimuli types. Dots are individual participants, the white dot reflects the median, the dark bar covers the interquartile range (IQR) with the whiskers indicating the maximum and minimum value within ±1.5 IQR. (B) Time-resolved power at the tagged frequencies in New trials. Colors indicate stimulus type. Shading indicates within-participant corrected standard error of the mean. (C) Response-locked power Old trials averaged over time, contrasting the stimuli types. Stars indicate significant differences. (D) Time-resolved power at the tagged frequencies in Old trials. Grey bar above plots indicates the significant cluster.

Next, we looked at visual search on Old trials, with established spatial predictive context. A priori time window analysis (Figure 4C) revealed stronger power for the Target than for distractors. Compared to D*near* this stronger Target power was evident in the data locked to the response (t(31) = 2.18, *p* = .019, *d* = .25, BF_10_ = 2.88), and marginally in the data locked to stimulus onset (t(31) = 1.66, *p* = .053, *d* = .16, BF_10_ = 1.23). Compared to D*far,* stronger power for the target was only present in the response-locked data (t(31) = 1.70, *p* = .049, *d* = .19, BF_10_ = 1.29), but not in stimulus-locked data (t(31) = .82, *p* = .21, *d* = .09, BF_10_ = .40). There was no difference between the distractor types (stimulus-locked: (t(31) = 1.10, *p* = .28, *d* = .14, BF_10_ = .33, response-locked: t(31) = .75, *p* = .46, *d* = .09, BF_10_ = .24).

Time-resolved analysis of Old trials (Figure 4D) further identified a positive cluster, especially comparing power at the target frequency to power at the near-standing distractor frequency (D*near*; cluster-based permutation test *p* = .018, *d* = .23, BF_10_ = 3.73). This cluster was identified in the early part of the response-locked window, from 550 to 250 ms before the response. There were no significant cluster-based differences between the target and the far-standing distractor, nor when contrasting the two distractor types in the response-locked window. There were additionally no significant clusters identified between any of the stimulus types during the stimulus-locked window.

We thus found evidence that spatial predictive context within the Old trials enhances the Target, compared to distractors. This biased competition appears strongest locally, between the target and the near-standing distractor specifically. However, we note that there was no direct evidence for a difference in power between the two distractor types.

### Target enhancement and distractor suppression in Old compared to New scenes bias local attentional competition

To see the direct impact of spatial predictive context on each stimulus type in isolation, we directly contrasted the power at each stimulus between Old and New scenes. We found evidence for higher target-related power in Old compared to New scenes, specifically when the data was aligned to the response (response-locked: t(31) = 2.32, *p* = .014, *d* = .15, BF_10_ = 3.75, stimulus-locked (t(31) = 1.06, *p* = .15, *d* = .09, BF_10_ = .53), but we found no evidence for lower distractor-related power in Old compared to New scenes (response-locked: D*near*: t(31) = .39, *p* = .35, *d* = .04, BF_10_ = .26, D*far*: t(31) = .13, *p* = .45, *d* = .01, BF_10_ = .21; stimulus-locked: D*near*: t(31) = 0.06, *p* = .48, *d* = .005, BF_10_ = .20, D*far*: t(31) = 0.14, *p* = .56, *d* = .015, BF_10_ = .17).

Next, we wanted to understand how the differences we find between Old and New trials interact with the attentional competition between stimuli during visual search. We therefore directly contrasted power during the a priori window on Old and New trials and subsequently compared the stimulus types, thus testing the interaction between stimulus type and condition. We found a difference between Target and both distractors when the data was aligned to the response (Target > D*near*: t(31) = 1.74, *p* = .046, *d* = .50, BF_10_ = 1.38; Target >D*far*: t(31) = 1.98, *p* = .028, *d* = .43, BF_10_ = 2.05), but not when data was aligned to the stimulus (Target > D*near*: t(31) = 0.84, *p* = .20, *d* = .22, BF_10_ = .41; Target > D*far*: t(31) = 0.86, *p* = .20, *d* = .17, BF_10_ = .42). There was no difference between the distractors (response-locked: t(31)= 0.15, *p* = .88, *d* = .04, BF_10_ = .19; stimulus-locked: t(31)= 0.14, *p* = .89, *d* = .04, BF_10_ = .19).

When analyzing these differences in attentional competition between Old and New trials in a time-resolved manner (Figure 5B), we found a significant cluster only when contrasting Target to D*near* (cluster-based permutation test *p* = .038, *d* = 0.59, BF_10_ = 3.16) from 350 ms to 450 ms before the response. There were no significant clusters when contrasting Target to D*far* or when contrasting the two distractor types, nor were there any significant differences during the stimulus-locked window. Visual inspection of figure 5A further supports these cluster based statistical results: target enhancement specifically co-occurs with distractor suppression of D*near*. The apparent time course of this effect explains why the cluster-based analysis reveals stronger evidence for this effect compared to the wider a priori defined time window (-0.8 to -0.2 s pre-response).

**Figure 5.**
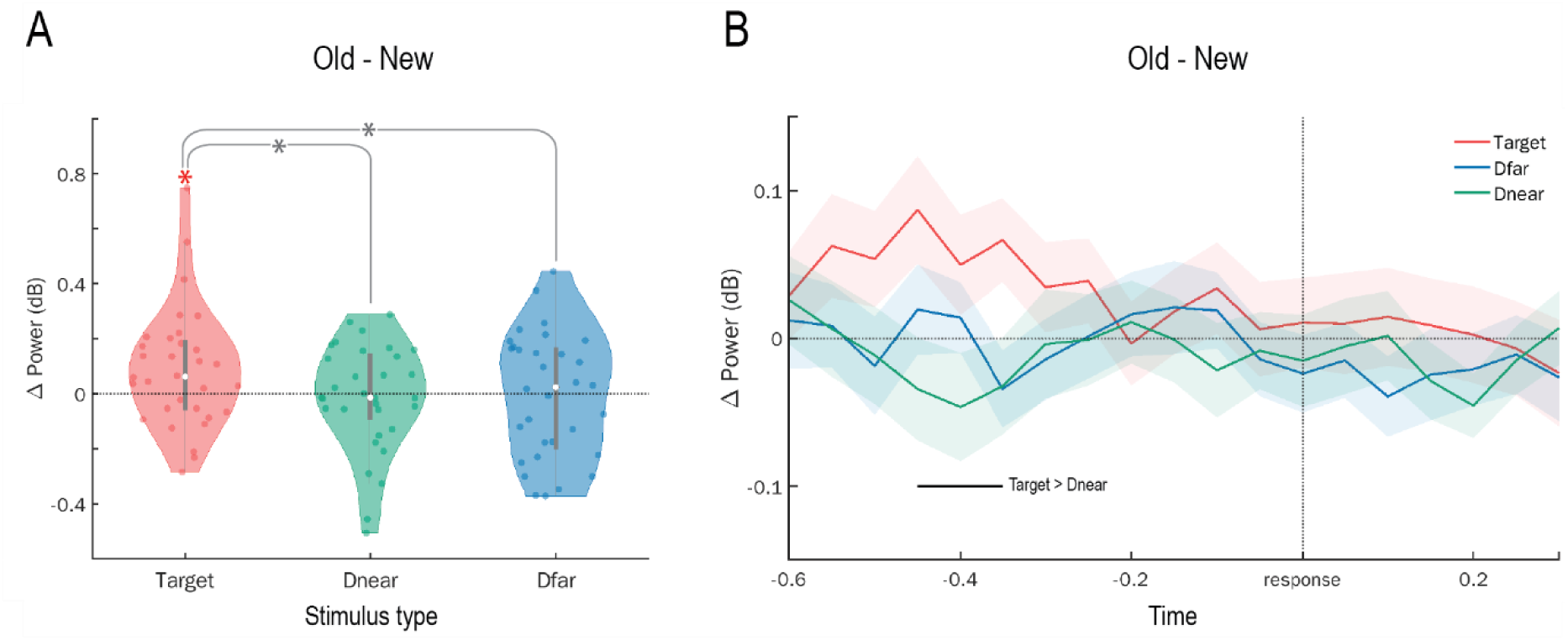
Response-locked power per stimulus type contrasting Old and New trials. (A) Response-locked power averaged across time, contrasting Old and New trials per stimulus type. Colored dots are individual participants, the white dot reflects the median, the dark bar covers the interquartile range (IQR) with the whiskers indicating the maximum and minimum value within ±1.5 IQR. Colored star indicates significant difference between Old and New trials. Grey stars indicate a significant difference of Old minus New power values between the stimuli types. (B) Time-resolved power. Shading indicates within-participant corrected standard error of the mean, the bar indicates the significant cluster.

Taking these results together, we conclude that the main difference between Old and New scenes is an enhancement of the target, and that this target enhancement specifically biases local attentional competition between the target and the directly surrounding distractors, which are subsequently suppressed.

### Both local and global competition are behaviorally relevant, but only local context is learned

Subsequently, we investigated whether this competition between target and distractors during visual search is meaningful for behavior. To do so, we tested whether an increase in power differences between target and distractors was related to an increase in search speed. We therefore correlated, within each subject, differences in power between these stimulus types with reaction times, across trials. To remove the overall effect of condition, we analyzed the New and Old trials separately (see Figure 6). By definition, aligning data to the time of the response removes any potentially interesting variance in reaction times and, therefore, a possible relationship between the MEG and reaction time data. Therefore, we analyze stimulus-locked data here.

**Figure 6.**
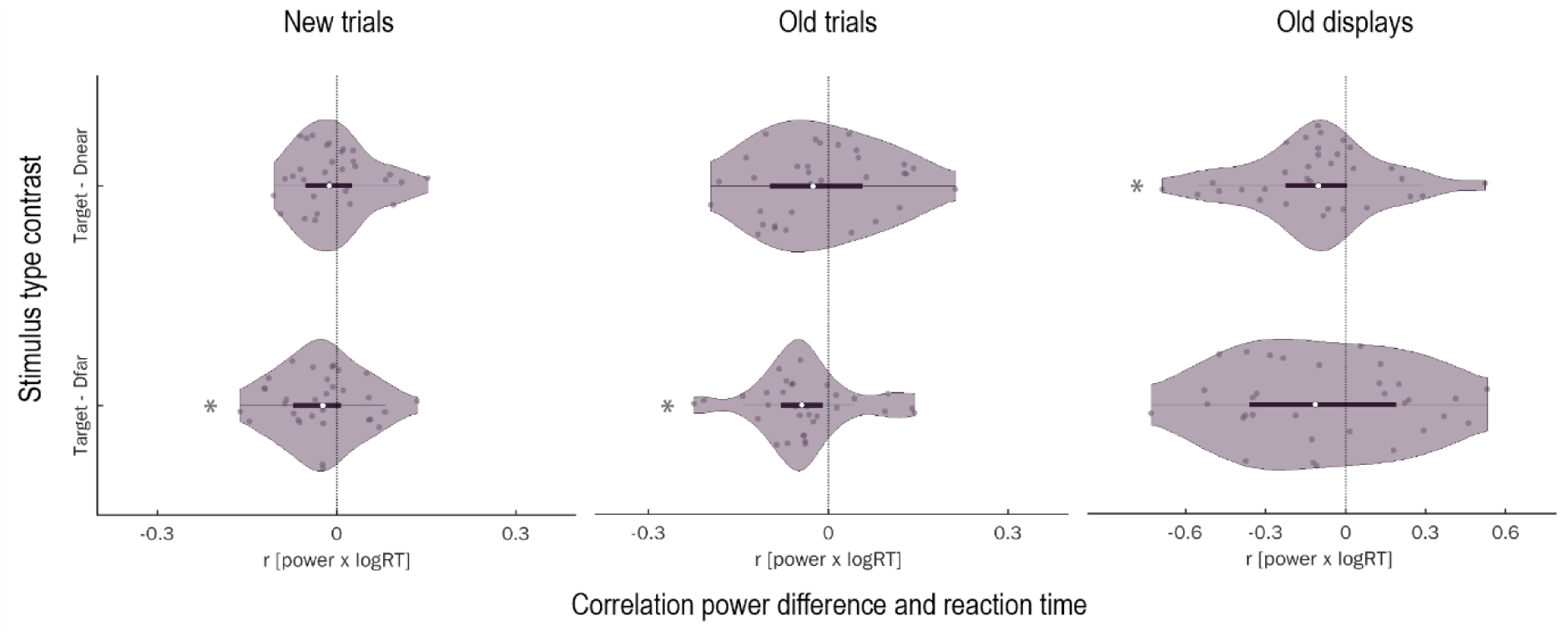
Behavioral relevance of stimulus-specific power. Distribution of r values expressing the relationship between stimulus-specific power and behavioral performance (logRT) per participant. Top plots contrast Target power to D*near* power, bottom plots contrast Target Power to D*far* power. From left to right the results for New trials, Old trials and Old trials averaged per display. Darker dots are individual participants, the white dot reflects the median and the grey bar indicates the interquartile range. Stars indicate that the r values are significantly different from zero.

On New trials, the difference between stimulus-locked Target power and power at D*far* was negatively correlated with reaction times, (mean *r* across participants = −.03, t(31) = 2.15, *p* = .020, *d* = .38, BF_10_ = 2.76), while the difference between Target and D*near* was not (mean *r* = −.005, t(31) = 0.44, *p* = .33, *d* = .08, BF_10_ = .27). We found a similar pattern within Old trials, with again the difference between Target and D*far* being predictive of search speed (mean *r* = −.04, t(31) = 2.68, *p* = .006, *d* = .47, BF_10_ = 7.71), and not the difference between Target and D*near* (mean *r* = −.01, t(31) = 0.67, *p* = .25, *d* = .12, BF_10_ = .35). We thus found a subtle, yet reliable effect of *global* competition: The difference in power between target and distractors is meaningful for general visual search performance, as participants were faster in finding the target when target power was stronger compared to power at far-standing distractors. This was true both for New and for Old trials.

We next sought to remove this general visual search effect, and isolate what impacts search performance on Old trials. These Old trials, by definition, consisted of repeating the same spatial configuration, or ‘display. This allowed us to remove general trial-by-trial variance and analyze the stimulus-locked power and reaction times averaged across Old trials per display (16 in total). Therefore, we averaged both the power differences and behavioral performance over the trials displaying the same (and thus repeated, Old) search scene, and calculated the correlation of the power differences and behavioral performance across these displays. We found that the power difference between Target and D*near* was correlated with behavioral performance (mean *r* = −.11, t(31) = 2.48, *p* = .009, *d* = .44, BF_10_ = 5.11), while this was not the case for the difference between Target power and D*far* (mean *r* = −.08, t(31) = 1.37, *p* = 0.089, *d* = .24, BF_10_ = .80). Thus, when target power was high compared to power at the near-standing distractor specifically, participants were faster in finding the target in Old scenes.

From this we conclude that, when there is more power for the target compared to distractors, people become faster in finding it. In general visual search, this holds specifically when target power is increased compared to distractors further away (presumably reflecting a general attentional drift effect, independent of learning). However, when we leverage the repetitions of Old displays, we are able to demonstrate the behavioral relevance of specifically competition between target and the near-standing distractor. This finding of behavioral relevance of *local* attentional competition between target and near-standing distractors dovetails with our results on stimulus-specific power both within Old trials separately, and in the contrast between Old and New.

### No evidence of explicit knowledge of Old scenes

*Recognition task*. With a mean accuracy of 51.46% (±11.05%), participants were at chance level in correctly identifying Old and New displays in the recognition task (Figure 7, t(31) = 0.75, *p* = .459, *d* = .13 BF_10_ = .25). Recognition accuracy was unrelated to whether participants were able to exploit predictive context behaviorally as we find no relationship between size of the contextual effect and recognition accuracy (r = 0.06, t(30) = 0.34, *p* = .734, BF_10_ = .41). We thus find no evidence of explicit knowledge of Old scenes, indicating that the learning visible in behavior was implicit.

**Figure 7.**
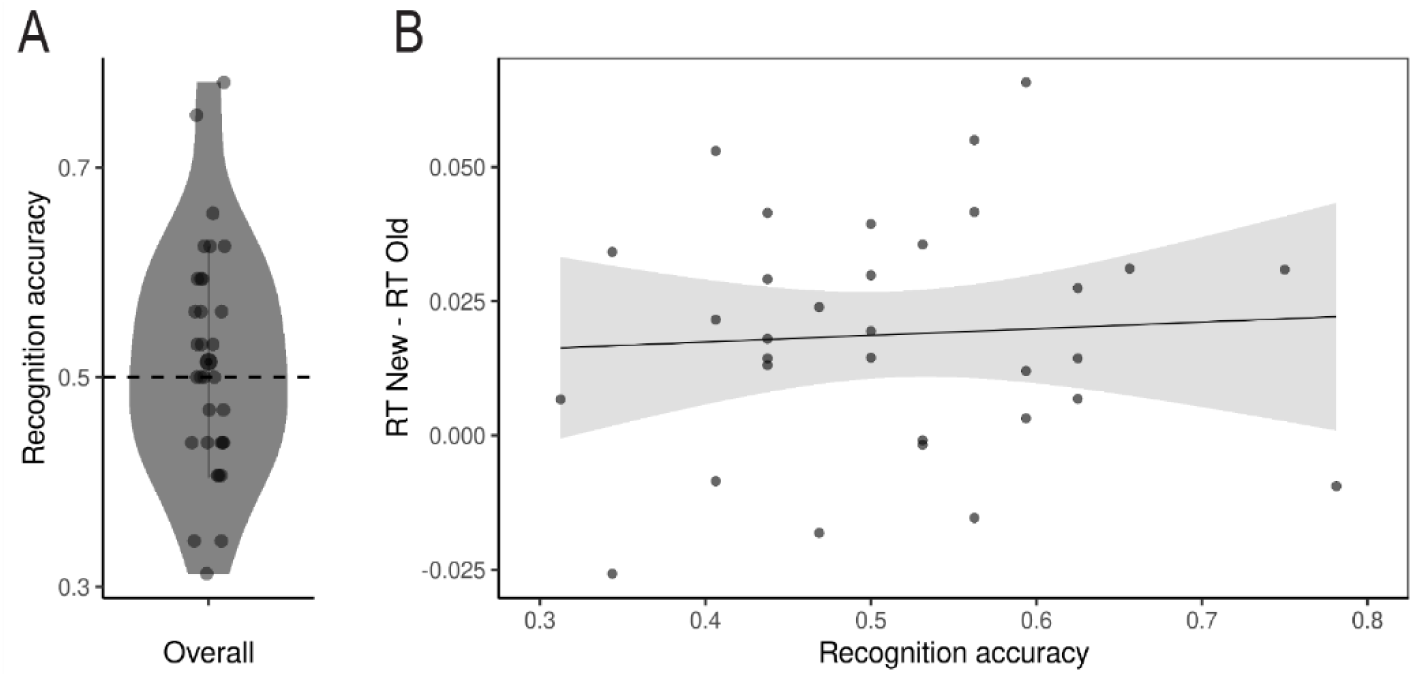
Recognition task. (A) Recognition accuracy (B) Contextual effect (RT New – RT Old) as a function of recognition accuracy. Dots in all panels represent individual participants.

## Discussion

Our study had two main objectives. Firstly, we aimed to investigate the neural mechanism underlying enhanced attentional processing of visual search when spatial context becomes predictive of target location. We utilized Rapid Invisible Frequency Tagging (RIFT) to track attentional processing of visual search scenes, and measure enhancement or suppression at the stimulus level. Secondly, by tagging two types of distractors, we probed the extent of the context involved in this improved attentional guidance. More specifically, we set out to test whether the entire context, including more distant distractors, or mainly the local context (i.e. distractors directly surrounding the target) play a role. Our findings revealed that visual search with predictive context (repeated scenes, referred to as “Old”), exhibited stronger Target-related power, especially compared to a distractor near that target. Moreover, contrasting predictive search to regular visual search without predictive context (referred to as “New”) reveals that both Target enhancement and distractor suppression underly this biased local attentional competition between the Target and directly surrounding distractors. Crucially, we found that for visual search in general, the difference between target and the far standing distractor was reflected in behavioral performance. However, when investigating Old displays specifically, we observed that biasing local competition between the target and the near standing distractor, is responsible for the accelerated search performance when spatial context is predictive. The implications of these findings are discussed below.

Our results indicate that the improved search performance in Old scenes is accompanied by stronger target power relative to distractors. This means that spatial predictive context creates a shift in shared attentional resources. More attention is allocated to the target, at the cost of the distractors, specifically those directly surrounding the Target. These results may explain several reported behavioral findings, such as contextual costs when the target in Old scenes is swapped with a distractor (Makovski & Jiang, 2010), and slower dot detection at learned distractor locations (Ogawa et al., 2007). Interestingly, our prediction would be that these impairments would only hold for local context and not for distractors further away, which has not been tested. Our findings additionally shed light on what underlies the enhanced N2pc found when there is spatial predictive context during visual search (Schankin & Schubö, 2009). The N2pc is an event-related potential (ERP) component believed to be related to attentional allocation. It relies on lateralized activity and therefore has limited granularity, and whether it reflects target selection, filtering out distractors, or both is under debate (Stoletniy et al., 2022). By leveraging RIFT, we were able to demonstrate that attentional allocation to the target within Old scenes is enhanced by biasing local competition in favor of the target, and at the cost of local distractors. Notably, Luck et al. (1997) demonstrated that increasing local competition enhances the N2pc. This finding is in line with our interpretation that spatial predictive context improves attentional guidance in visual search by biasing this local competition.

Moreover, the comparison between Old and New scenes revealed a simultaneous occurrence of target enhancement and distractor suppression. Previous research has implicated both in visual search, also in more natural settings (Hickey et al., 2009; Seidl et al., 2012). It has been argued that these processes rely on different mechanisms (Hickey et al., 2009; Noonan et al., 2016; Slagter et al., 2016; Wöstmann et al., 2022). However, distractor suppression has been mainly established using exogenous or otherwise obvious cues manipulating the location probability of a target or distractor (e.g., cues indicating distractor location, distractor consistently in the same hemifield, etc.) that can induce strategic and therefore ‘proactive’ distractor suppression (Ferrante et al., 2023; Wöstmann et al., 2022). Here, using a contextual cueing paradigm, we demonstrate that distractor suppression can also operate when strategic suppression based on explicit knowledge is highly unlikely. However, we did not establish distractor suppression in isolation: we found no differences between Old and New trials when looking at either distractor. Distractor suppression is known to be weaker compared to target enhancement (Wöstmann et al., 2022). We furthermore only measure two distractors (one far, one near the target), while suppression is expected to be shared between multiple distractors (including those not tagged). When contrasting the distractors to the target, we find that both target enhancement and distractor suppression are jointly biasing attentional competition towards the target, at the cost of specifically local distractors. We demonstrate a tight coupling between target enhancement and distractor suppression during visual search, indicative of an attentional field with surround suppression (Hopf et al., 2006; Müller et al., 2005; Störmer & Alvarez, 2014). This coupling of target enhancement and distractor suppression, specifically locally, is indicative target-context associative learning. However, future research is needed to understand the neural consequences of context learning independently of the target, and how this contributes to contextual cueing (Beesley et al., 2015a; Vadillo et al., 2021).

We found that the repetition of search scenes primarily affects the processing of spatial context directly surrounding the target. This finding supports the notion that local context plays a critical role in contextual cueing (Brady & Chun, 2007; Goujon et al., 2015; Sisk et al., 2019). Previous research has demonstrated that merely repeating the local context is sufficient to produce similar contextual cueing to repeating the entire spatial array (Brady & Chun, 2007). However, this mainly indicates that the visual system will exploit what can be learned from our environment, it does not prove global context to be redundant. In our study, we directly demonstrate the importance of local context, even when global context is available. We speculate that the relationship between the attentional field and learning in contextual cueing is bi-directional: the attentional field strengthens the learning of local context and this learned local context subsequently sharpens the attentional field, ultimately sidelining global context. This implies a need for sufficient time to process the local context around the target during visual search, before this bi-directional effect can come into play. Interestingly, Xie et al. (2020) demonstrated that only repeating local context does *not* yield the same contextual cueing effect as repeating the entire search array when presentation time is very short. This bi-directional learning would explain why different behavioral contextual cueing studies have found that local context is still linked to the global configuration (Brady & Chun, 2007; Y. Jiang & Wagner, 2004; Zheng & Pollmann, 2019). Local context is local only because it is surrounded by global context, since using fewer stimuli actually diminishes the contextual cueing effect (Kunar et al., 2007). If global context guides attention towards the target, we anticipated this modulation early in the search process, as it should bias the searcher to the correct part of the scene, and this would go at the cost of Dfar which is placed in the opposite (xy) position. In our results we were not able to find such evidence of global context impacting configural learning. We believe this mostly stresses the relative importance of local context but does not exclude the possibility that global context might still be important. Further research is needed to fully understand the role of global context as it clearly plays a role in natural scenes that evoke more global processing (Brockmole et al., 2006).

Importantly, we were able to demonstrate the behavioral relevance of target power compared to the far-standing distractor on all *trials*, both Old and New. This may serve as evidence of efficient versus inefficient visual search (Schlagbauer et al., 2017; Tseng & Li, 2004), where higher power at the target relative to the distractor at the opposite position of the visual field indicates that the random starting point of search was immediately in the right direction. Our analyses of Old *displays*, instead, show the behavioral advantage of Target power compared to the near standing distractor. This means that spatial predictive context does not simply speed up visual search by biasing towards the target, but fundamentally changes the process by biasing local attentional competition.

The time course of biased local attentional competition during visual search with spatial predictive context reveals additional aspects. First, when aligning to stimulus onset we did not observe any apparent differences between Old and New trials. This suggests that very early differences between Old and New scenes (Chaumon et al., 2008) are not responsible for the behavioral advantage. Instead, we did find a difference between Old and New trials in our data aligned to the response, reaching its maximum at approximately 400 ms pre-response and waning well before the response. The response-locked nature of our results is compatible with the notion that contextual cueing facilitates response selection (Kunar et al., 2007, 2008). However, it is also compatible with an account of contextual cueing enhancing attentional guidance predominantly in the later part of the search process. (Kunar et al., 2007, 2008). Interestingly, a recent eye-tracking study found the benefit of predictive context during visual search consistently during the final 500 ms of search, independent of set size (Harris & Remington, 2020). The number of fixations until the target was found was reduced for Old displays, a signature of improved attentional guidance, but this reduction was found later in the search process when the set size was larger. The putative explanation for this consistently late enhancement is the importance of local context in contextual cueing (Harris & Remington, 2020; Sisk et al., 2019). We now demonstrate that indeed local competition in a time window close to the end of the search process is relevant for when there is spatial predictive context.

It is of importance to note that our experimental design subtly deviated from the ‘classic’ contextual cueing design. In the original setup from Jiang & Chun, targets on New and Old trials had separate locations. We did not have this restriction in our design, and this makes our results less generalizable to existing contextual cueing literature. Our design inevitably led to some target locations occurring more frequently than others, and this could potentially have led to ‘target probability cueing’. A recent interesting study by Geyer et al. (2024), showed that sharing target locations between Old and New trials led to location probability learning on New trials and, by consequence, a reduced contextual cueing effect. Our control analyses showed no evidence of target probability learning on New trials. We can, however, not exclude the possibility that target probability learning is present on Old trials. It is possible that spatial (distractor) context and target location were learned separately, in the extreme case without any association being formed between them. However, as spatial context and target location were always co-occurring on Old trials, and given the highly established nature of contextual cueing, across a variety of experimental designs, we deem it plausible that an association was formed between them. However, we stress that we cannot rule out alternative scenarios, such as the one in which Old trials form a more reliable context in which target probability learning may occur (Geyer et al., 2024). In any case, the neural results we report are of interest under any of the interpretations of the exact mechanisms underlying the type of contextual learning at play here.

Our results advance the technique of RIFT in several ways. Previous RIFT studies have predominantly involved low attentional competition, with spatial attention often being explicitly cued to a hemifield (Seijdel et al., 2023; Zhigalov et al., 2019) and a maximum of four, typically large, stimuli on the screen (Ferrante et al., 2023). In our visual search paradigm, stimuli were considerably smaller, spatial attention was more distributed, and competition was uncued, resulting in much subtler differences between stimuli. The fact that we were able to measure attentional competition and the biasing thereof by spatial predictive context highlights the strength of this innovative method.

Lastly, as expected, we established contextual cueing in participants’ behavior, demonstrating a robust and reliable difference wherein they were faster in locating a target on Old trials compared to New trials. Notably, participants performed at chance level when tested on their memory of these Old scenes. Moreover, participants who demonstrated larger contextual cueing benefits, did not have higher accuracy rate in the recognition task. Therefore, we conclude that our results provide no evidence of explicit knowledge acquisition of spatial predictive context.

In conclusion, our findings provide compelling evidence that spatial predictive context improves attentional guidance during visual search. It does so by enhancing the target and suppressing directly surrounding distractors, sharpening the attentional field. This leads to biased attentional competition enabling improved search performance. These results thus further our understanding of how humans can quickly and implicitly learn from their environment and exploit this learning to their benefit.

## Acknowledgements

We would like to thank Jorie van Haren for his help with collecting the data.

